# PI4KIIIβ inhibition reduces rhinovirus associated cell shedding and ciliary dysfunction

**DOI:** 10.1101/2023.12.14.571686

**Authors:** Simona A. Velkova, Alina M. Petris, Dani D.H. Lee, Daniela Cardinale, Dale Moulding, Richard A. Williamson, Soren Beinke, Ken Grace, Edith M Hessel, Nikolai N. Belyaev, Tanja Hoegg, Michael Steiner, John R Hurst, Rosalind L. Smyth, Claire M. Smith, Primrose Freestone, Christopher O’Callaghan

**Affiliations:** Infection, Immunity and Inflammation Research & Teaching Department, UCL Great Ormond Street Institute of Child Health, 30 Guilford Street, London, WC1N 1EH, United Kingdom; Department of Respiratory Sciences, University of Leicester, Leicester, LE1 9HN, United Kingdom; GSK Medicines Research Centre, Gunnels Wood Road, Stevenage, SG1 2NY, United Kingdom; Development Biology and Cancer Program, UCL Great Ormond Street Institute of Child Health, 30 Guilford Street, London, WC1N 1EH, United Kingdom; Institute for Lung Health, University of Leicester, LE3 9QP, Groby Road, Leicester, United Kingdom; UCL Respiratory, Rayne Building, University College London, WC1E 6JF, United Kingdom; Eligo Bioscience, 29 Rue du Faubourg Saint-Jacques Paris, 75014 France

## Abstract

**Background:** Patients with chronic obstructive pulmonary disease (COPD) experience respiratory exacerbations, many of which are associated with rhinoviruses. Current treatment strategies do not target the pathogenic rhinovirus trigger.

**Research question:** What is the immediate effect of rhinovirus on the ciliated respiratory epithelium and can viral replication and epithelial toxicity be reduced by targeted PI4KIIIβ inhibition.

**Methods:** Short (24h) and longer (7 days) rhinovirus infection were explored in primary ciliated airway epithelial cultures from multiple healthy and COPD patients using high-speed video microscopy, viral titration assays and immunofluorescence studies. Ciliated epithelial cultures were pre-treated with a PI4KIIIβ (GSK’533) blocker prior to infection to assess efficacy against rhinovirus. Cytokine and chemokine production were assessed by multiplex immunoassays.

**Results:** Within hours of infection rhinovirus co-localised with ciliated cells causing extensive apoptosis-associated shedding of predominantly ciliated cells within 24 hours. Viral replication that peaked at day 1 and cleared by day 7, was associated with dramatic loss of ciliated cells confirmed by reduced ciliary activity and ciliary DNAI2 protein expression. Ciliary beat frequency (CBF) of remaining cilia was significantly reduced by day 7 in cultures from COPD. Infection was partly dependent on PI4KIIIβ with the GSK’533 blocker reducing viral replication while preserving ciliary activity. High levels of pro-inflammatory mediators were secreted by infected cells.

**Conclusion:** Decreased ciliation due to rhinovirus infection is likely to impair mucociliary clearance in healthy individuals and COPD patients, contributing to the pathophysiology of COPD exacerbations. PI4KIIIβ inhibition blocks viral replication, helping to preserve ciliary activity.

**Take home message:** Rhinovirus replication in the healthy and COPD respiratory epithelium is mediated by PI4KIIIβ and intracellular PI4P platform formation. Inhibition of PI4KIIIβ reduced viral replication and ciliated cell loss.

## Introduction

Rhinoviruses account for almost half of colds in adults and a quarter of colds in children [1, 2]. Infection typically causes mild upper respiratory tract symptoms in healthy individuals but is a common trigger of wheezing in young children and patients with asthma [3]. Rhinovirus causes 60% of viral respiratory exacerbations in COPD patients and can cause severe and prolonged exacerbations [4–7]. Rhinovirus is thought to spread directly from person to person through infectious respiratory secretions or indirectly through contaminated surfaces [5, 8], however, rhinovirus detection in exhaled breath [9] suggests a potential role of aerosol transmission. Despite the huge clinical impact of rhinovirus, no vaccine and no specific antiviral treatment are currently available, and current interventions to treat and prevent exacerbations of COPD are incompletely effective.

Human rhinoviruses are single stranded RNA positive viruses from the *Picornaviridae* family of Enteroviruses, with more than 160 known serotypes [3]. The rhinovirus genome consists of approximately 7200 base pairs and it replicates rapidly yielding around 100,000 virions per cell within six hours [10]. Epithelial cells are recognised as the main site of rhinovirus infection [3, 8, 11] with ciliated cells being targeted for replication [12, 13].

The current study focuses on rhinovirus 16 which is thought to bind to intracellular adhesion molecule-1 (ICAM-1) receptors present on the plasma membrane of airway epithelial cells, although of interest, the ICAM-1 receptor has not been found on ciliated or squamous epithelial cells [1, 8, 14]. PI4KIIIβ kinase regulates membrane transport from the trans-Golgi network to the plasma membrane via PI4P (phosphatidylinositol-4-phosphate) [15, 16]. It has been reported that picornaviridae (the family to which rhinoviruses belong) and flaviviruses hijack PI4KIIIβ enzymes and PI4P lipids for the formation of replication complexes in infected cells [17–19]. The majority of research into mechanisms of rhinovirus infection have involved immortalised epithelial cell lines, which may not reflect *in vivo* infection [11, 20]. The aim of this study was therefore, to investigate infection of ciliated primary respiratory epithelial cells, grown at an air fluid interface from multiple COPD patients and healthy individuals to help reflect variability in response, and to more closely mimic *in vivo* infection. We also evaluated the effect of inhibition of PI4KIIIβ [16] on rhinovirus infection.

### Methodology

An expanded version of the methods can be found in Supplementary Materials.

### Collection of primary human respiratory epithelial cells

Primary airway epithelial cells were collected by nasal brushings [21, 22]. Patient details are summarised in Supplementary Table 1. Ethical approval for sample collection was obtained through the Living Airway Biobank (Ref: 14/NW/0128).

### Culture of primary human respiratory epithelial cells

Co-culture of human airway cells with mitotically inactivated feeder layers was carried out as previously described [22, 23]. Human primary cells were cultured at 4x10^5^cells/0.33cm^2^ on transwell insert (Corning, USA) in submerged conditions for 2 days at 37°C and 5%CO_2_. Thereafter, primary cells were grown at air-liquid interface (ALI) and transepithelial electrical resistance (TEER) was measured as previously described [21, 22].

### Virus culture

Rhinovirus 16 provided by Prof Gary McLean (Cellular and Molecular Immunology Research Centre, London, UK) was propagated in Hela H1 cells [24] and infected cell lysates were purified and titred [25].

### Rhinovirus *in vitro* infection of human airway ciliated cultures and PI4KIII**β** inhibition assay

An aliquot of 1.5x10^6^TCID_50_/ml was added to the apical site of epithelial cultures in 100μl BEBM media and removed after 1h. Mock infected cultures received 100μl BEBM media and were processed as infected wells. In inhibition assays, ALI cultures were pre-treated for 1h with 100nm of PI4KIIIβ inhibitor (GSK’533) [16] or vehicle control (DMSO) basolaterally prior viral infection.

### Ciliary function analyses

Recordings of cilia beating were carried out using a Hamamatsu ORCA digital camera C11440 (Hamamatsu, Japan) attached to an inverted phase-contrast Nikon Eclipse Ti-E (Nikon instruments, UK) microscope and a 20x long distance objective at 394 frames per second (fps) at 512x512 pixel resolution with 2ms exposure time. Ciliary beat frequency (CBF) and ciliary beat activity were determined using the ciliaFa software [26].

### Immunofluorescence microscopy

Epithelia culture samples fixed in 4% (w/v) PFA were stained in transwell inserts [22, 27] and imaged by Zeiss LSM 710 confocal microscope (Carl Zeiss ltd, Germany). Primary antibodies anti-VP2 (QED biosciences, USA), anti-βtubulin (Abcam, UK), anti-PI4P (Echelon Biosciences, UK) were detected by anti-rabbit A488 (Abcam, UK), anti-mouse A647 (Biorad, UK) and anti-mouse A488 (Abcam, UK). Anti-VP2 antibody was also conjugated to A647 fluorophore (Abcam, UK) to avoid host species cross-reactivity. Hoechst 33258 (Sigma-Aldrich, UK) was applied to counterstain cell nuclei.

### Western blot analysis

Western blots were carried out using standard procedures [22]. Antibodies anti-DNAI2 (1:500, mouse, Bio-Techne) and anti-GAPDH (1:2000, rabbit, Abcam) were detected by anti-mouse (Biо-techne) and anti-rabbit (Cell signalling technologies) horseradish peroxidase linked antibodies. Densitometry analysis was performed using ImageJ software 1.51j8 (National Institutes of Health, USA). To determine the abundance of the target proteins, percent values were normalised to relative GAPDH.

### Apoptosis analysis

Epithelial cells detached from the epithelial layer post infection were stained with UV zombie dye (BioLegend) and Annexin V dye (BD Pharmingen) and fixed in 1%PFA. Cells were cytospun onto glass slides. Images were taken on a Nikon TiE-Eclipse 100 microscope using a 100x objective.

### Cytokine and chemokine measurement

Cytokines and chemokines were measured using 96-well MSD plates (Meso Scale, UK) and custom Luminex plex plates (Invitrogen, UK) as per manufacturers’ instructions.

### Statistical analysis

Statistical analysis was done in R (R-core team 2018) (see supplementary methods), Version 3.5.1 with Benjamini-Hochberg multiple comparison correction. Graphs were plotted using GraphPad Prism v6.1 (GraphPad software, USA).

## Results

### Rhinovirus rapidly targets and detaches ciliated epithelial cells during the first day of infection

Six hours post infection immunofluorescent staining showed co-localisation of rhinovirus with ciliated cells (70%) (Figure 1A). Staining was intensive in ciliated cells, suggesting so much greater replication in this cell type. Light microscopy and high-speed video imaging showed epithelial shedding of ciliated cells started as early as 6 hours post infection, which continued over the 24h study period (Figure 1B shows a single frame from a video that can be seen in Supplementary Video 1). Loss of ciliated cells was supported by reduced ciliary beat activity; 24h post infection there was a 71% decrease with estimated relative change 0.29 [95% Cl: 0.087 – 0.934], (p=0.042) in healthy and 66% decrease with estimated relative change 0.34 [95% Cl: 0.104 – 1.108], (p=0.064) in COPD compared to mock-infected cultures (Figure1C). When ciliated cells were shed from the epithelium, their shape quickly changed from columnar to a rounded phenotype (Figure 1B). Cilia on detached cells continued to beat for up to 24 hours in an increasingly dyskinetic fashion before becoming static. Epithelial cultures remained intact with no difference in TEER values between infected and non-infected cultures (Figure 1E).

**Figure 1:**
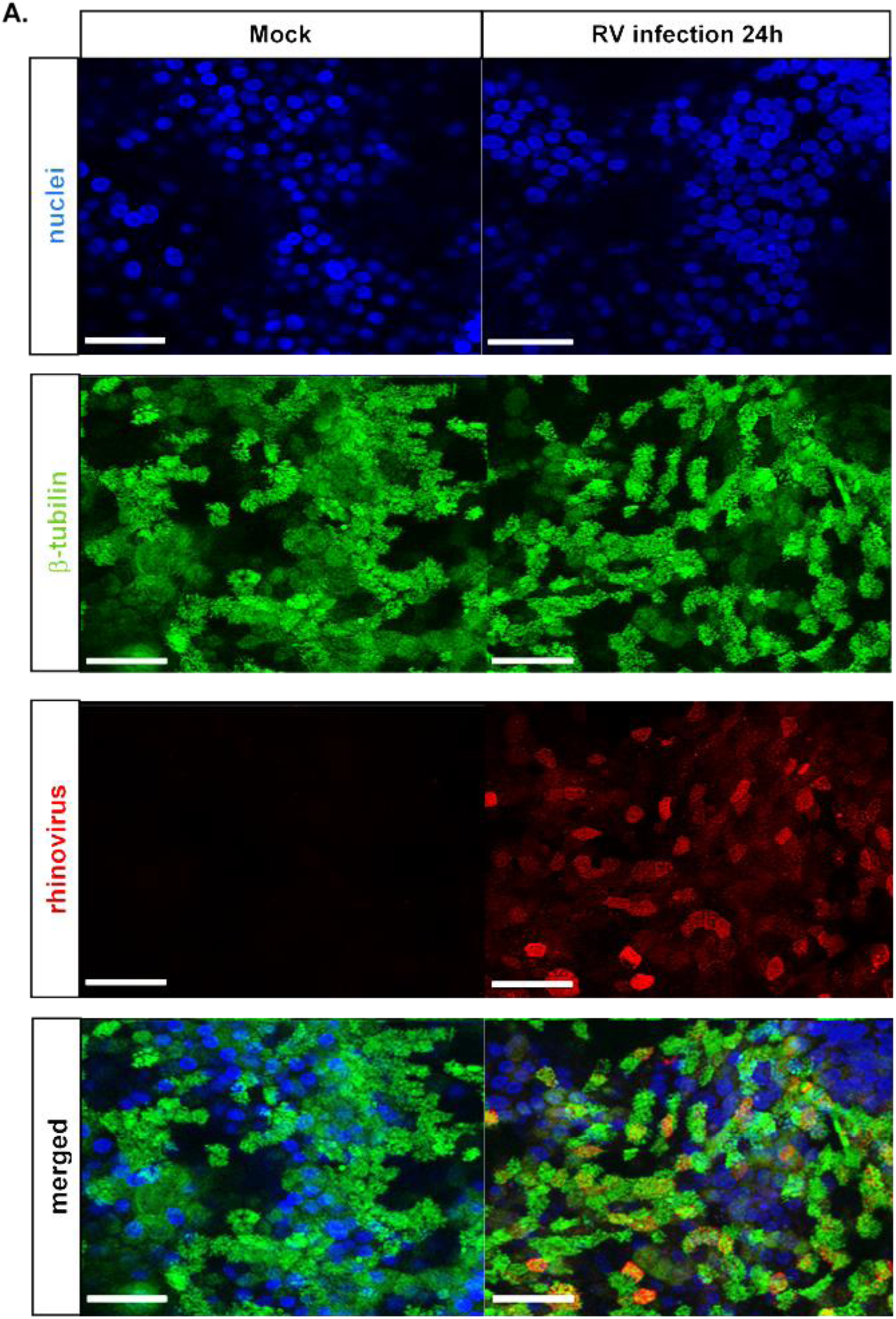
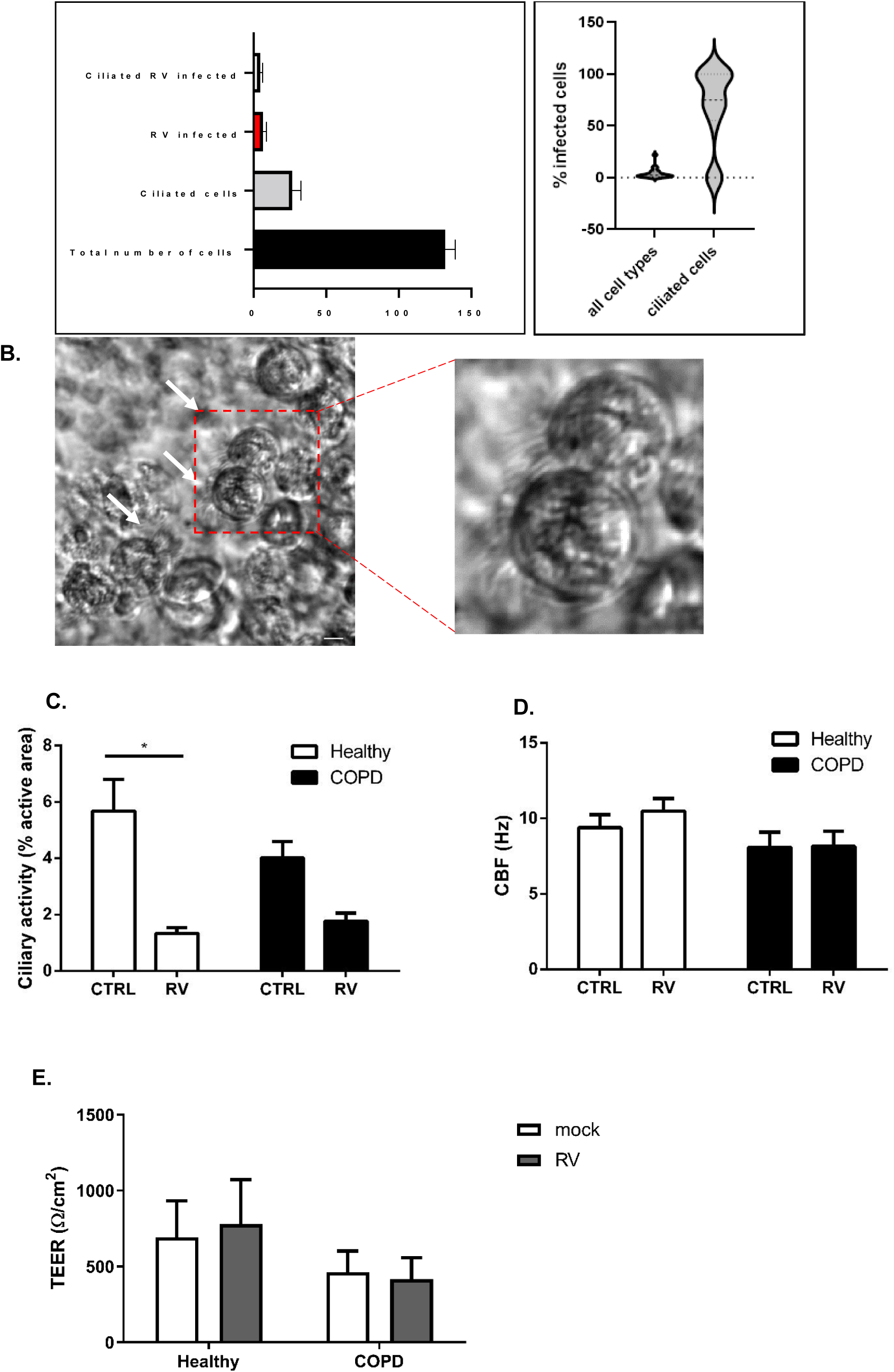
Rhinovirus infection of the airway ciliated epithelium at early time-points. A) The confocal images show that at 6 hours post infection: rhinovirus was strongly associated with ciliated cells. Cilia were stained with anti-β-tubulin antibody (green), rhinovirus with anti-VP2 antibody (red), nuclei with Hoechst solution (blue) and imaged by LSM710 Zeiss confocal microscope, scale bars= 20μm; graphs show mean number of cells (DAPI+/ β-tubulin+/ RV VP1+) counted per image (n=7 images/donor), 2 Healthy and 1 COPD donor and extrapolated percent infected cells at 6h; B) is a screenshot of a COPD epithelium infected with rhinovirus, imaged at 18 hour post infection by high speed video microscopy, white arrows point at detached ciliated cells, scale bar = 10μm; C) Ciliary activity of ciliated cultures expressed as % of the area where beating cilia were detected following rhinovirus infection, n=10 areas/donor, n=3/group; D) Ciliary beat frequency (CBF) of ciliated cultures following rhinovirus infection, n=10 areas/ donor; E) TEER of epithelial cultures at 24h post RV infection, 6 Healthy and 6 COPD donors, error bars represent mean ± SEM., *=p<0.05;

CBF of cilia present on cells still attached to the epithelial layer at 24hours post infection, was measured in 10 pre-selected epithelial fields of view (Figure 1D). No significant difference in CBF was seen between rhinovirus-infected cultures from healthy donors (10.5 ± 0.8Hz) compared to mock-infected cultures (9.4 ± 0.9Hz, p=0.2) or from infected cultures from COPD donors (8.2 ± 1Hz) and mock-infected cultures (8.1 ± 1Hz, p=0.89) (Figure 1D). No significant difference in baseline CBF was found in healthy and COPD cultures (p=0.47).

During this experiment there was no significant difference between the amount of RV in apical fluid at 24hour between healthy (5.5x10^4^ ± 2.6x10^4^ TCID_50_/ml, n=6) and COPD donors (3.3x10^4^ ± 1.4x10^4^ TCID_50_/ml, n=6, p=0.56). In the absence of apical fluid there was a non-significant increase of viral titre in COPD (3.2x10^4^ ± 1.6x10^4^ TCID_50_/ml, n=6) compared to healthy (2.4x10^4^ ± 4x10^3^ TCID_50_/ml, n=6, p=0.56) (data not shown).

### Effect of rhinovirus on epithelial cytokine and chemokine production

Cytokine and chemokine levels were measured in apical and basolateral fluids to investigate response to rhinovirus infection. The measurements results are summarised in Supplementary Tables 2&3.

Our data shows a significant increase in a number of mediators associated with recruitment of inflammatory cells measured 24hours after infection of COPD cultures in comparison to mock-infected cultures (Figure 2A, Supplementary Tables 2&3). Cytokines IL-1β and IL-6 (p<0.05), IP-10, RANTES and IFNλ2/3 (p<0.0001) all increased. In cultures from healthy donors, IL-6 (p<0.01), IP-10, RANTES and IFNλ2/3 (p<0.0001) increased significantly. In our infection model IL-17c, a potent TH17 immune cell stimulator in COPD [28], was detected significantly higher post rhinovirus infection in COPD and in healthy controls (p<0.0001). GM-CSF was statistically increased apically in cultures from COPD donors (p<0.05), but basolaterally in healthy donors (p<0.001) (Figure 2A, 2C). The macrophage and neutrophil chemotactic chemokines IL-8 (p<0.05 healthy, COPD), MIP-3α (p<0.01 healthy, p<0.05 COPD), MCP-1 (p<0.001 healthy, p<0.01 COPD) and TNF-α (p<0.01 healthy) were significantly higher in the basolateral fluids of infected cultures (Figure 2A). We also report for the first time that IL-15, an important cytokine in T cell and natural killer cells activation [29], was significantly increased in rhinovirus COPD secretions (p<0.01)(Figure 2C). Additionally, chemokines ENA-78, TARC, GRO-α, CXCL14 were non-significantly elevated post rhinovirus infection (Supplementary Table 2). Principal component analysis showed rhinovirus infection in COPD shifts the inflammatory mediator response in the basolateral site of cultures with main contributors TARC, IFN-β and IL17c (Figure 2B), suggesting heightened COPD response for T cells and specifically Th17. Interestingly, inflammatory response of COPD to rhinovirus on the apical site was deficient of interferons (IFNα, IFNβ, IFNλ) in contrast to non-infected and healthy controls (Figure 2D).

**Figure 2:**
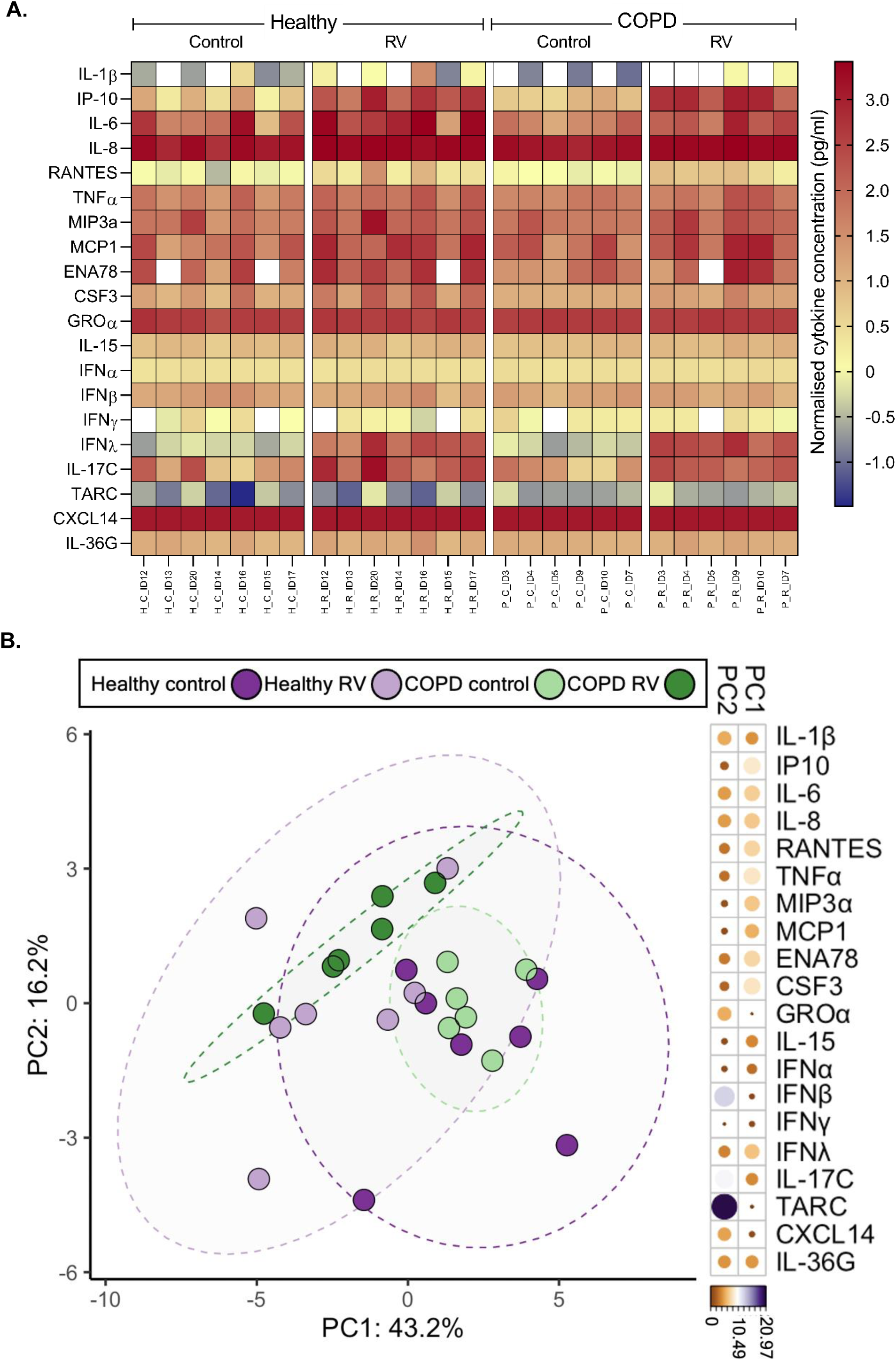
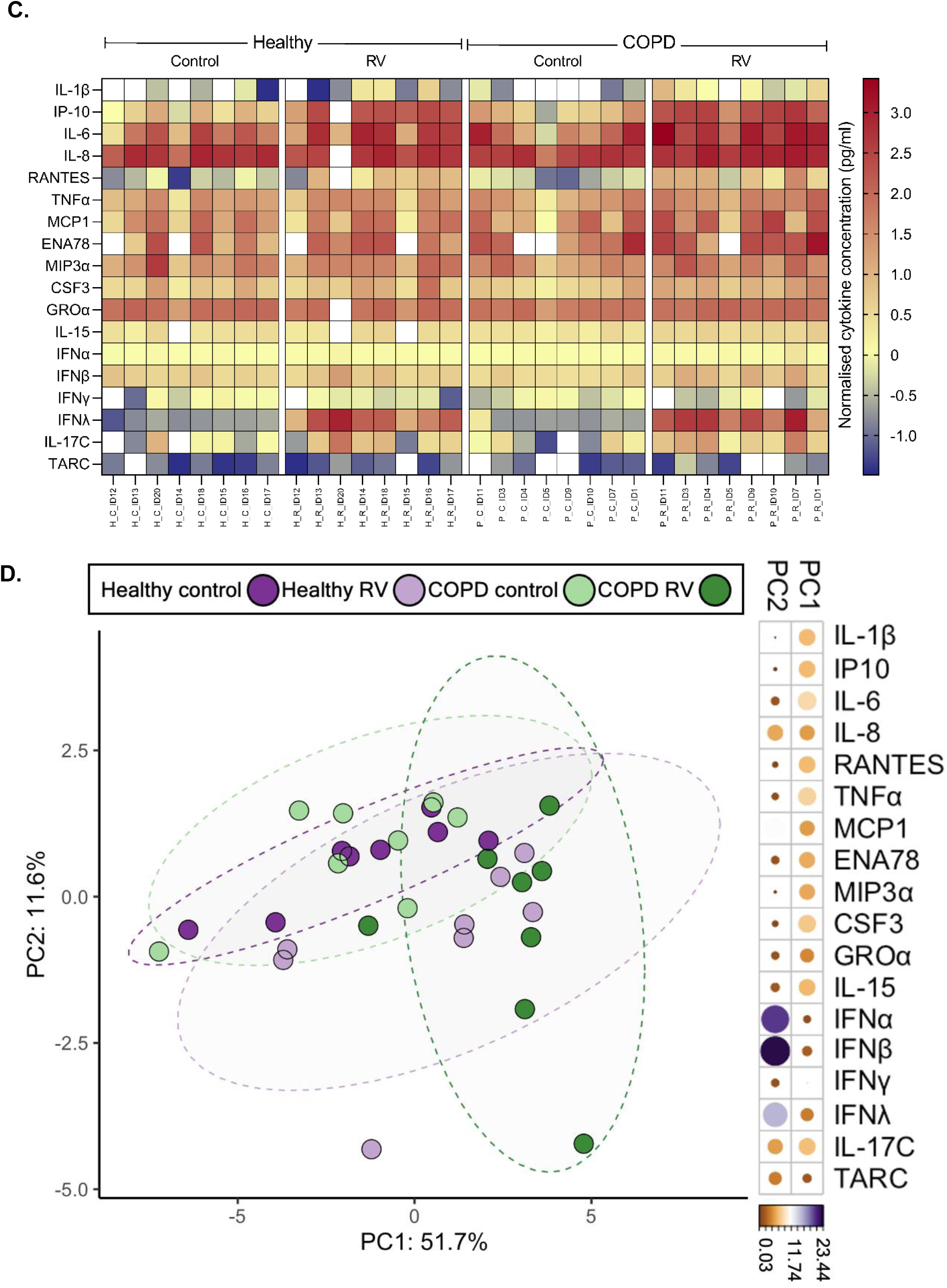
Inflammatory mediators secreted by the airway ciliated epithelium at 24h post rhinovirus infection. Inflammatory mediators secreted in the basolateral fluids of epithelial cultures from healthy or COPD donors infected with rhinovirus for 24h. (A) Heat map of inflammatory mediators secreted basolaterally; (B) PCA showing variation of response to rhinovirus from epithelial cultures originating from healthy and COPD donors, n=7 donors. Inflammatory mediators secreted in the apical fluids of epithelial cultures from healthy or COPD donors infected with rhinovirus for 24h. (C) Heat map of inflammatory mediators secreted apically; (D) PCA showing variation of response to rhinovirus from epithelial cultures originating from healthy and COPD donors, n=7 donors

### Ciliated cells are lost and viral replication eliminated by 7 days rhinovirus infection of the respiratory epithelium

Ciliary activity of 4 healthy and 4 COPD donor cultures was studied for 7 days post infection, and was reduced compared to mock infection in all donors at all time points (Figure 3A). In addition to reduction of ciliary activity, there was a visual reduction of ciliated cells within 24hours of infection with further loss increasing over time. Reduced DNAI2 expression (a quantitative marker of cilia presence) was seen with rhinovirus infection compared to mock infection (Figure 3B). Our results are consistent with significant cilia loss, suggesting reduced ciliary activity was due to loss of ciliated cells rather than ciliary stasis. Supplementary Figure 1 shows Western blots of reduced DNAI2 protein expression. Day 7 HSV microscopy recordings were taken on a daily basis to analyse cellular dynamics and intactness of the epithelial layer (Supplementary Figure 2C). Previous experience show damage to epithelium presents with flooding but this was not seen.

**Figure 3:**
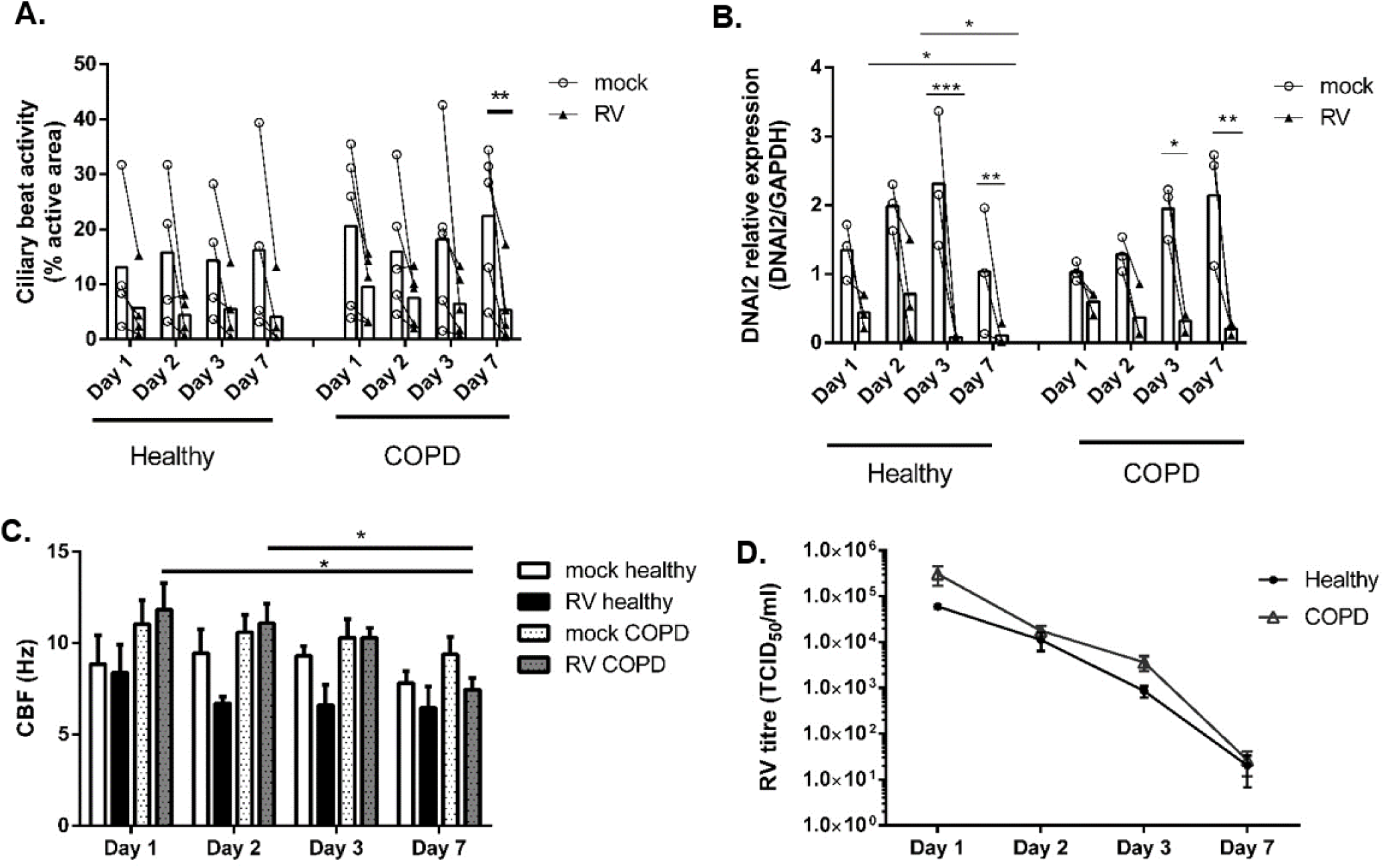
Long-term (7 day) rhinovirus infection of the airway ciliated epithelium. A) Ciliary activity in mock and rhinovirus infected ciliated cultures at 1, 2, 3 and 7 days post rhinovirus infection: n=4/group with lines following individual donors; B) DNAI2 expression relative to GAPDH expression in mock and rhinovirus infected ciliated cultures at days 1, 2, 3 and 7 post rhinovirus infection: n=3/group with lines following individual donors; C) Ciliary beat frequency (CBF) (Hz) of mock and rhinovirus infected nasal ciliated cultures at days 1, 2, 3 and 7 post infection, n=10 field of views/ donor with n=4 donors/group n=4/group, error bars represent mean±SEM; D) Live rhinovirus titre in nasal airway cultures at days 1, 2, 3 and 7 post rhinovirus infection, n=4/ group, error bars represent ±SEM; *=p<0.05, **=p<0.01, ***=p<0.001.

Figure 3C shows maintenance of CBF of cilia on ciliated cells that remained attached to the respiratory epithelium. A significant reduction of CBF was seen in COPD at day 7 of infection in comparison to days 1 (-4.3Hz [95% Cl:-6.6Hz – -2.1Hz], p=0.01) and 2 (-3.6Hz [95% Cl:-5.9Hz – -1.3Hz], p=0.04).

Again, at 24hours there was a non-significant increase of viral titre in cultures from COPD donors (3x10^5^ ± 1.4x10^5^ TCID_50_/ml) compared to healthy (6x10^4^ ± 5 x10^3^ TCID_50_/ml, p=0.18) (Figure 3D). As the infection progressed, viral load gradually decreased. A significant reduction of replicating live virus, compared to day 1, was found in cultures from healthy and COPD individuals, at days 2 (p≤0.01), 3 (p<0.001) and 7 (p<0.001). Viral load in ciliated cultures was mostly eliminated by day 7, in cultures from healthy (2.8x10^2^ ± 1.6x10^2^ TCID_50_/ml) and COPD donors (2.7x10^1^ ± 1.5x10^1^ TCID_50_/ml). Our findings suggest rhinovirus viral replication peaks at 24hours and is almost undetectable by day 7 with the ciliated respiratory epithelium, in the absence of immune cells, playing a pivotal role in the resolution of rhinovirus infection. A non-significant reduction in TEER (transepithelial electrical resistance) was seen on day 7 of infection (60% reduction with estimated relative change of 0.4 [95% Cl: 0.15-1], p=0.26 in cultures from healthy donors and 51% reduction with estimated relative change of 0.49 [95% Cl: 0.17-1.42], p=0.38 in cultures from COPD donors) (Supplementary Figure 2).

### Ciliated epithelial cell shedding occurs via apoptosis

As described earlier (Figure 1) rapid shedding of cells from the epithelium was observed following rhinovirus infection (Figure 4). We studied detached epithelial cells in apical secretions of ciliated cultures from 4 healthy and 4 COPD donors in 25 pre-selected areas/donor with standard surface of 0.015mm^2^/ per viewing area. Figure 4A represents the total number of cells and total number of ciliated cells detached from the epithelium. In healthy donors, apical fluids of mock infected cultures contained only detached cells with cilia (mean% ± SEM %): 3% ± 2%, compared with rhinovirus infected cultures from healthy donors: 38% ± 8%. In COPD, again, apical fluids of mock infected cultures contained 6% ± 2% detached cells with cilia in contrast to 24% ± 7% in rhinovirus infected cultures.

**Figure 4:**
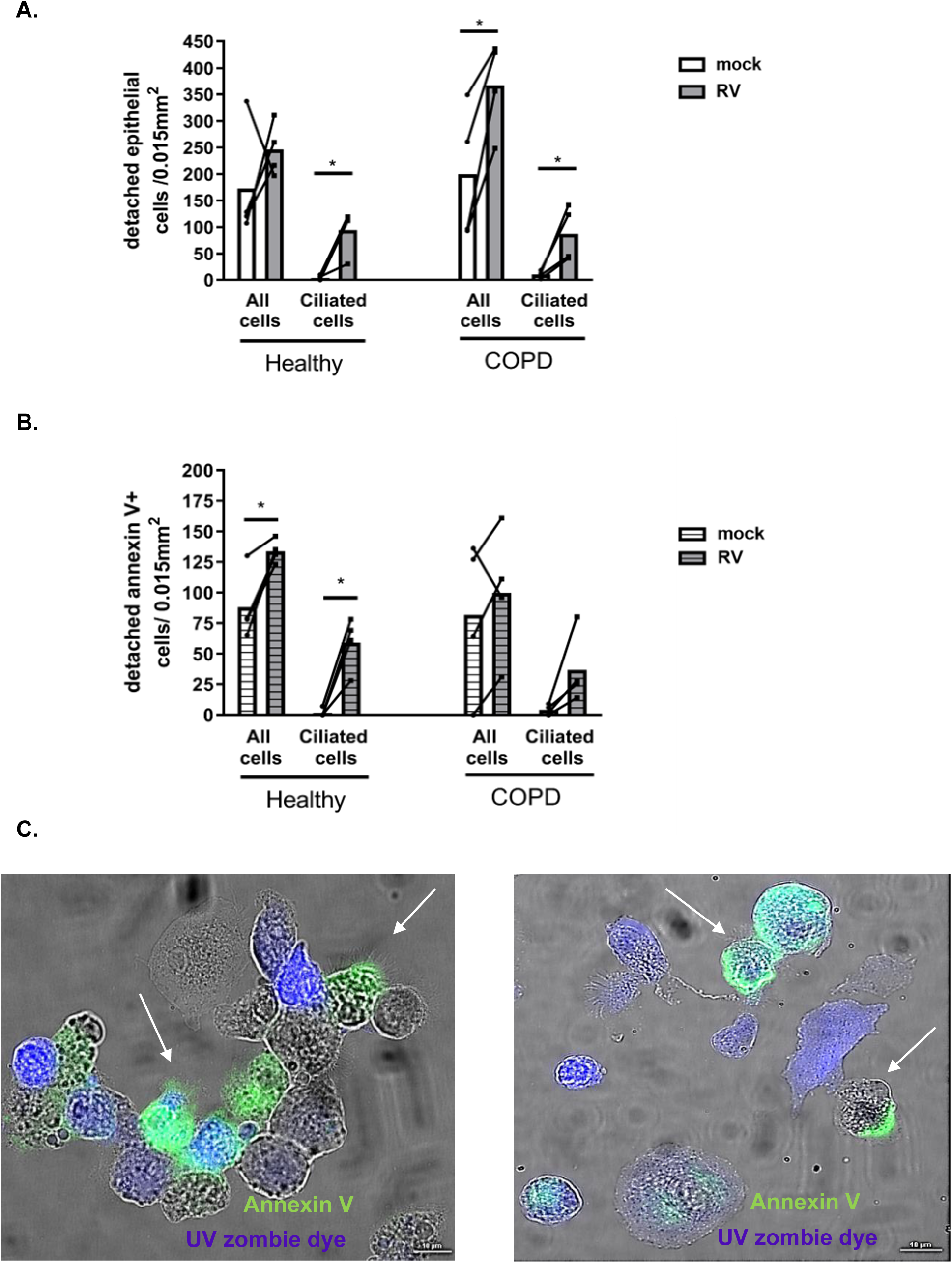
Programmed cell death in rhinovirus infected airway ciliated cultures at 24h. A) Number of total cells and ciliated cells detached from the epithelium, collected by apical washing; B) Number of total cells and ciliated cells detached from the epithelium, collected by apical washing, positive for Annexin V; Cells in A) and B) were counted per 0.015mm^2^ field from 25 random images using 100x objective, lines follow individual donors, n=4 donors/ group; C) Representative images of epithelial cells in the apical washes of cultures following rhinovirus infection showing ciliated cells positive for apoptotic cell markers. Apoptotic (Annexin V+) cells appear in green, late apoptotic (Annexin V+; UV zombie dye+) cells appear in green and blue, necrotic/ dead (UV zombie dye+) cells appear in blue, scale bars=10μm. *=p<0.05

Apical secretions were stained with Annexin V, an early apoptotic marker and UV zombie dye, a necrosis marker (Figure 4B and C). In cultures from healthy donors, in mock infected secretions, (mean% ± SEM%): 1%±1% of cells with cilia were Annexin V positive in contrast to 24%±5% in rhinovirus infected secretions. Similarly, in COPD, in mock infected secretions, only 1%±1% of cells with cilia were Annexin V positive compared to 37%±6% in rhinovirus infected secretions. Actual numbers are presented in Figure 4B. Of interest, very few necrotic cells found in apical secretions after rhinovirus infection were ciliated (Supplementary Figure 3).

### Blocking PI4KIIIβ enzyme activity maintains ciliary activity

To investigate whether PI4KIIIβ kinase activity was involved in rhinovirus replication in differentiated human airway epithelium and if it correlated with cell shedding, we pre-treated mucociliary cultures with a PI4KIIIβ inhibitor (GSK’533) [16, 30]. Inhibition of PI4KIIIβ in cultures significantly reduced viral replication (Figure 5A). 100nM of GSK’533 was enough to significantly decrease the titre of live virus in ciliated cultures from healthy and COPD (p<0.0001) donors compared to vehicle controls. On observation there was a clear reduction of ciliated cells shed from the PI4KIIIβ treated rhinovirus infected epithelium compared to untreated rhinovirus only infection. Ciliary activity was measured to determine if blocking PI4KIIIβ enzyme activity prior to infection preserved ciliated cells. In all but one of the donor cultures, a similar trend of reduced ciliary activity was seen post rhinovirus infection in contrast to the mock and PI4KIIIβ inhibitor treated cultures where ciliary activity was maintained. Treatment with GSK’533 significantly preserved ciliary activity (p=0.047) of rhinovirus infected cultures from healthy individuals. The trend was similar in two COPD cultures but not in the third (Figure 5B). The PI4KIIIβ blocking agent did not alter CBF (Figure 5C) or TEER readings (Figure 5D).

**Figure 5:**
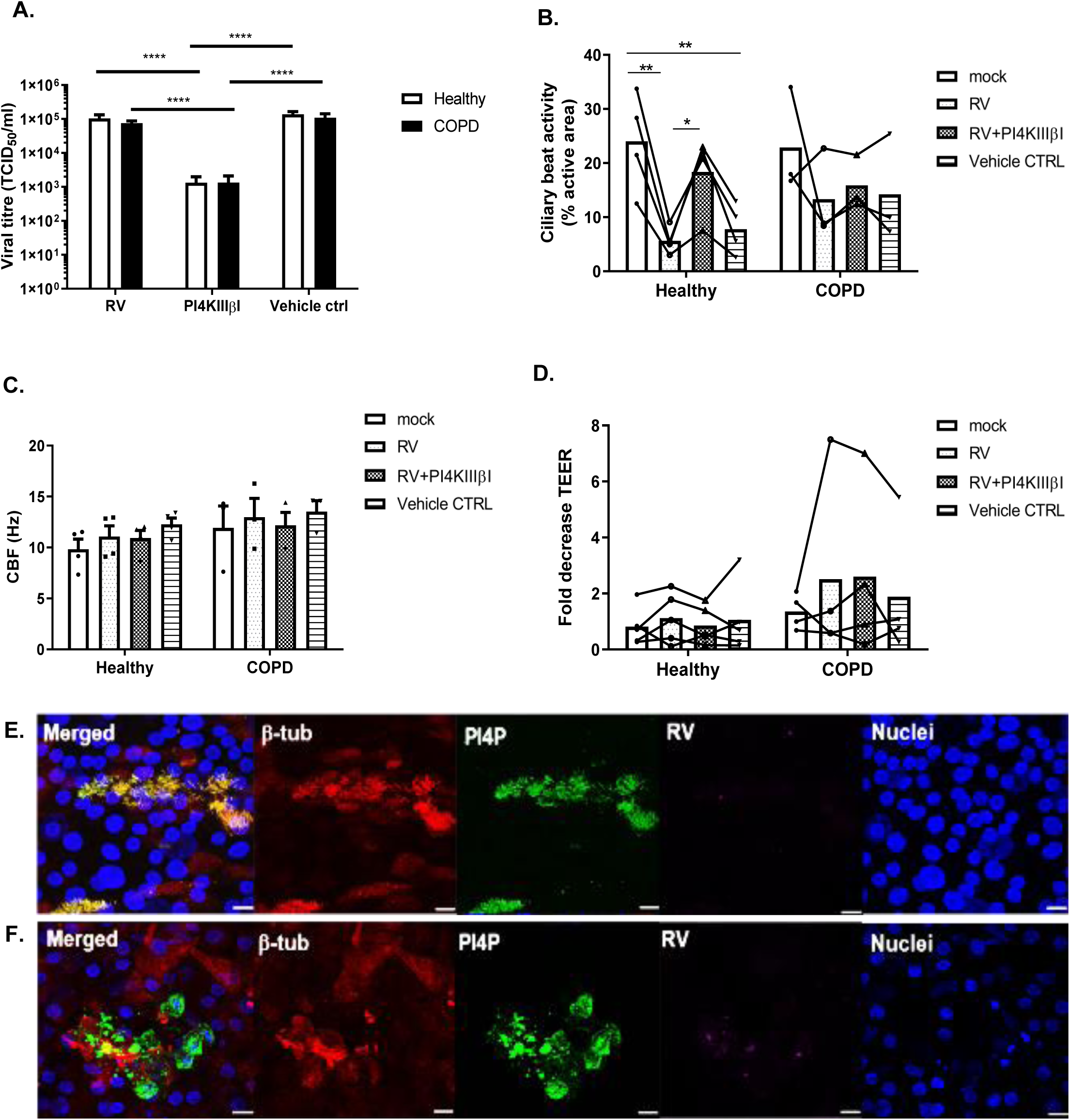
PI4KIIIβ inhibition decreases rhinovirus replication and associated cell shedding in airway ciliated cultures 24hour post infection. A) Live rhinovirus titre in rhinovirus-infected airway ciliated cultures untreated, in presence of PI4KIIIβ inhibitor or vehicle control, n=4/ group; B) Ciliary beat activity of ciliated cultures in rhinovirus-infected airway ciliated cultures untreated, in presence of PI4KIIIβ inhibitor or vehicle control column bars represent mean values, n=10 fields of view for 4 healthy donors, 3 COPD donors with lines following individual donors; C) Ciliary beat frequency (CBF) of ciliated cultures in rhinovirus-infected airway ciliated cultures untreated, in presence of PI4KIIIβ inhibitor or vehicle control, n=10 fields of view for 4 healthy donors, 3 COPD donors; D) Transepithelial electrical resistance (TEER) of nasal ciliated cultures in rhinovirus-infected airway ciliated cultures untreated, in presence of PI4KIIIβ inhibitor or vehicle control, relative to pre-treatment readings, n=2 in 4 donors/ group, lines following individual donors; E) Positive green fluorescent PI4P staining is coincident with red fluorescent β-tubulin staining in non-infected control, confirming PI4P appearance in motile cilia; F) Positive green fluorescent PI4P staining is no longer seen in cilia but inside epithelial cells, where it overlaps with β-tubulin in red and rhinovirus in magenta colour staining; Scale bars = 16μm; Error bars represent mean±SEM, *=p<0.05, **=p<0.01, ***=p<0.001, ****=p<0.0001

To confirm the presence of PI4P in respiratory epithelial cells, fully differentiated cultures were stained with an anti-PI4P antibody. In mock infected cultures (Figure 5E) a high expression of PI4P was seen in motile cilia, along the entire length of the ciliary axoneme. PI4P expression was also visualised intracellularly where expression was clear but much lower. Infected cultures (Figure 5F) experiencing extensive ciliated cell loss, showed a marked and enriched expression of PI4P intracellularly. These findings appeared similar in cultures from healthy and COPD donors.

## Discussion

The aim of this study was to examine the effect of rhinovirus on the ciliated respiratory epithelium and investigate if rhinovirus replication and disruption to ciliary function was reduced by targeted PI4KIIIβ inhibition. We demonstrated in ciliated cultures from multiple healthy and COPD donors that early viral interaction with the airway epithelium was characterised by tropism for ciliated cells, with marked shedding of predominantly ciliated cells within 24hours of infection, with preservation of epithelial integrity. The majority of cells shed from the epithelium were apoptotic with a small number of necrotic cells seen. It is unclear whether shedding of infected cells from the epithelium with apoptosis benefits the host by clearing the infection or the virus by enhancing potential spread through epithelial secretions, or indeed both.

Our results suggest the characteristic nasal symptoms of the common cold, rhinorrhoea and nasal obstruction, experienced shortly after rhinovirus infection is significantly contributed to by the marked reduction in ciliation of respiratory epithelia, reducing mucociliary clearance. Of interest, in the absence of inflammatory cells, the cultured ciliated epithelium was still able to clear rhinovirus infection with no virus detectable after seven days. This finding is consistent with studies showing viral elimination of respiratory syncytial virus (RSV) [31] and parainfluenza [32] from cell cultures by day 8 post infection.

Confocal imaging suggested tropism of rhinovirus for ciliated cells leading to their shedding, consistent with other reports [12, 33, 34]. However, we report here for the first time that rhinovirus infection particularly reduces CBF in cultures from COPD patients over time. The combination of profound deciliation of the epithelium combined with defective ciliary function in COPD is likely to reduce mucociliary clearance contributing to clinical exacerbations if replicated in the lower airway. If mimicked in small airways rapid cellular extrusion may also contribute to airway obstruction. Although our study involved nasal epithelial cells a similar degree of loss of ciliary activity and CBF was seen in bronchial culture from a COPD patient following rhinovirus infection (data not shown). Loss of ciliated epithelia could also provide a predisposing mechanism for recurrent bacterial exacerbation which has been reported following initial viral events [4], and an explanation for the funding that exacerbations tend to cluster in time with a high-risk period for a recurrent event in the period following [35].

This study also determined that PIKIIIβ inhibition in primary ciliated cultures from COPD patients and healthy individuals reduced rhinovirus replication and epithelial disruption. Our investigation showed abundant PI4P expression in cilia as opposed to the cell bodies prior to infection. Following rhinovirus infection that was associated with marked ciliary depletion, enriched intracellular PI4P expression was seen. PIKIIIβ inhibition, in cultures from healthy and COPD donors, significantly reduced rhinovirus replication and ciliated cell detachment from the epithelium, suggesting PIKIIIβ involvement in the pathophysiology of rhinovirus infection as previously proposed [16, 30]. The phosphoinositides (PI) present in the plasma membrane of many cell types in different tissues, are involved in cellular signalling and cascades as second messengers [36]. PI4P from the PI lipid family is synthesised by PI4KIIIβ and PI4KIIα kinases, and localised to the cell Golgi apparatus with major role to recruit factors for Golgi-endosomal vesicular transport [15]. It was recently reported RNA viruses can utilise PI4K enzymes, enhancing the production of PI4P lipids at the Golgi network and together with viral replication proteins and host cells molecules, build machinery for efficient viral synthesis and proliferation [17, 19]. The PI4KIIIβ enzyme was implicated in cell lines as a vital contributor to viral platform formation for viruses including SARS coronavirus, poliovirus, coxsackievirus, hepatitis C [17, 37].

Rhinovirus infection elicited a strong inflammatory response in ciliated cultures from healthy and COPD donors. Principal component analysis revealed rhinovirus heightened COPD response for T cells driven by TARC, IL17c and IFNβ, likely influencing the pathogenesis of disease. Consistent with our results IL-17c was recently found increased in the sputum of COPD patients during exacerbations and was linked to disease severity [38]. Here, we provide novel *ex vivo* data that the IL-15 protein was significantly elevated in the apical surface of cultures from COPD donors following rhinovirus infection. Recently, in an *in vivo* mouse study, rhinovirus stimulated type Ι IFN dependent IL-15 protein expression to the apical surface of the airway epithelial lining where it was required for natural killer and CD8+ T cells recruitment and activation [29]. In an *in vivo* influenza infection study, IL-15 presented on pulmonary dendritic cells promoted virus specific CD8+ T cells survival and lung accumulation [39], however the relevance of IL-15 in the pathogenesis of rhinovirus infection in COPD remains unclear.

We found significantly higher levels of IL-8, MIP-3α and MCP-1, known to be chemotactic to macrophages and neutrophils, in the basolateral fluids of infected cultures. IL-1β, IL-6, IP-10, RANTES were significantly increased in the apical and basolateral fluids of airway cultures from healthy and COPD donors following infection with no significant differences between the two groups. Our findings are consistent with previous reports of these cytokines found in COPD sputum, BAL samples and in *in vitro* culture viral infections [3, 4, 8, 40].

In conclusion, we have shown rhinovirus rapidly infects ciliated cells of primary airway epithelial cultures from multiple healthy and COPD donors. Infection led to shedding of airway cells from the differentiated epithelium, with a dramatic reduction of ciliation within 24hours and induction of apoptotic cell death. Viral infection resulted in a strong pro-inflammatory release of cytokines from the epithelium. Pre-treatment of cultures from different donors with a PI4KIIIβ inhibitor reversed rhinovirus replication and cell detachment from the epithelium, preserving ciliation. Thus, PI4KIIIβ inhibition may provide a therapeutic target for rhinovirus infection, helping to preserve mucociliary clearance in healthy individuals and during COPD exacerbations. Better treatment and prevention strategies for COPD exacerbations are urgently required, especially those that target RV. We found ciliated respiratory epithelial cultures are able to clear rhinovirus replication within 7 days suggesting inflammatory cells *in vivo* may not be essential for viral elimination.

## Supporting information

Supplementary Material - Results

## References

1. Bella, J. and M.G. Rossmann, Review: rhinoviruses and their ICAM receptors. J Struct Biol, 1999. 128(1): p. 69–74.

2. van der Zalm, M.M., et al., Highly frequent infections with human rhinovirus in healthy young children: a longitudinal cohort study. J Clin Virol, 2011. 52(4): p. 317–20.

3. Bochkov, Y.A. and J.E. Gern, Rhinoviruses and Their Receptors: Implications for Allergic Disease. Curr Allergy Asthma Rep, 2016. 16(4): p. 30.

4. George, S.N., et al., Human rhinovirus infection during naturally occurring COPD exacerbations. Eur Respir J, 2014. 44(1): p. 87–96.

5. Wedzicha, J.A. and G.C. Donaldson, Exacerbations of chronic obstructive pulmonary disease. Respir Care, 2003. 48(12): p. 1204–13; discussion 1213-5.

6. Seemungal, T., et al., Respiratory viruses, symptoms, and inflammatory markers in acute exacerbations and stable chronic obstructive pulmonary disease. Am J Respir Crit Care Med, 2001. 164(9): p. 1618–23.

7. Mallia, P., et al., Experimental rhinovirus infection as a human model of chronic obstructive pulmonary disease exacerbation. Am J Respir Crit Care Med, 2011. 183(6): p. 734–42.

8. Jacobs, S.E., et al., Human rhinoviruses. Clin Microbiol Rev, 2013. 26(1): p. 135–62.

9. Leung, N.H.L., et al., Author Correction: Respiratory virus shedding in exhaled breath and efficacy of face masks. Nat Med, 2020. 26(6): p. 981.

10. Johnston, S.L., P.G. Bardin, and P.K. Pattemore, Viruses as precipitants of asthma symptoms. III. Rhinoviruses: molecular biology and prospects for future intervention. Clin Exp Allergy, 1993. 23(4): p. 237–46.

11. Lopez-Souza, N., et al., Resistance of differentiated human airway epithelium to infection by rhinovirus. Am J Physiol Lung Cell Mol Physiol, 2004. 286(2): p. L373–81.

12. Griggs, T.F., et al., Rhinovirus C targets ciliated airway epithelial cells. Respir Res, 2017. 18(1): p. 84.

13. Tan, K.S., et al., In Vitro Model of Fully Differentiated Human Nasal Epithelial Cells Infected With Rhinovirus Reveals Epithelium-Initiated Immune Responses. J Infect Dis, 2018. 217(6): p. 906–915.

14. Winther, B., et al., Surface expression of intercellular adhesion molecule 1 on epithelial cells in the human adenoid. J Infect Dis, 1997. 176(2): p. 523–5.

15. Tokuda, E., et al., Phosphatidylinositol 4-phosphate in the Golgi apparatus regulates cell-cell adhesion and invasive cell migration in human breast cancer. Cancer Res, 2014. 74(11): p. 3054–66.

16. Roulin, P.S., et al., Rhinovirus uses a phosphatidylinositol 4-phosphate/cholesterol counter-current for the formation of replication compartments at the ER-Golgi interface. Cell Host Microbe, 2014. 16(5): p. 677–90.

17. Altan-Bonnet, N. and T. Balla, Phosphatidylinositol 4-kinases: hostages harnessed to build panviral replication platforms. Trends Biochem Sci, 2012. 37(7): p. 293–302.

18. Ilnytska, O., et al., Enteroviruses harness the cellular endocytic machinery to remodel the host cell cholesterol landscape for effective viral replication. Cell Host Microbe, 2013. 14(3): p. 281–93.

19. Hsu, N.Y., et al., Viral reorganization of the secretory pathway generates distinct organelles for RNA replication. Cell, 2010. 141(5): p. 799–811.

20. Edwards, M.R., et al., New treatment regimes for virus-induced exacerbations of asthma. Pulm Pharmacol Ther, 2006. 19(5): p. 320–34.

21. Hirst, R.A., et al., Ciliated air-liquid cultures as an aid to diagnostic testing of primary ciliary dyskinesia. Chest, 2010. 138(6): p. 1441–7.

22. Lee, D.D.H., et al., Ciliated Epithelial Cell Differentiation at Air-Liquid Interface Using Commercially Available Culture Media. Methods Mol Biol, 2020. 2109: p. 275–291.

23. Butler, C.R., et al., Rapid Expansion of Human Epithelial Stem Cells Suitable for Airway Tissue Engineering. Am J Respir Crit Care Med, 2016. 194(2): p. 156–68.

24. Abo-Zeid, Y., et al., An investigation of rhinovirus infection on cellular uptake of poly (glycerol-adipate) nanoparticles. Int J Pharm, 2020. 589: p. 119826.

25. Bartlett, N.W., A. Singanayagam, and S.L. Johnston, Mouse models of rhinovirus infection and airways disease. Methods Mol Biol, 2015. 1221: p. 181–8.

26. Smith, C.M., et al., ciliaFA: a research tool for automated, high-throughput measurement of ciliary beat frequency using freely available software. Cilia, 2012. 1:p. 14.

27. Smith, C.M., et al., Ciliary dyskinesia is an early feature of respiratory syncytial virus infection. Eur Respir J, 2014. 43(2): p. 485–96.

28. Pfeifer, P., et al., IL-17C is a mediator of respiratory epithelial innate immune response. Am J Respir Cell Mol Biol, 2013. 48(4): p. 415–21.

29. Jayaraman, A., et al., IL-15 complexes induce NK- and T-cell responses independent of type I IFN signaling during rhinovirus infection. Mucosal Immunol, 2014. 7(5): p. 1151–64.

30. Spickler, C., et al., Phosphatidylinositol 4-kinase III beta is essential for replication of human rhinovirus and its inhibition causes a lethal phenotype in vivo. Antimicrob Agents Chemother, 2013. 57(7): p. 3358–68.

31. Liesman, R.M., et al., RSV-encoded NS2 promotes epithelial cell shedding and distal airway obstruction. J Clin Invest, 2014. 124(5): p. 2219–33.

32. Zhang, L., et al., Infection of ciliated cells by human parainfluenza virus type 3 in an in vitro model of human airway epithelium. J Virol, 2005. 79(2): p. 1113–24.

33. Mosser, A.G., et al., Similar frequency of rhinovirus-infectible cells in upper and lower airway epithelium. J Infect Dis, 2002. 185(6): p. 734–43.

34. Jakiela, B., et al., Th2-type cytokine-induced mucus metaplasia decreases susceptibility of human bronchial epithelium to rhinovirus infection. Am J Respir Cell Mol Biol, 2014. 51(2): p. 229–41.

35. Hurst, J.R., et al., Temporal clustering of exacerbations in chronic obstructive pulmonary disease. Am J Respir Crit Care Med, 2009. 179(5): p. 369–74.

36. Winkler, D.G., et al., PI3K-delta and PI3K-gamma inhibition by IPI-145 abrogates immune responses and suppresses activity in autoimmune and inflammatory disease models. Chem Biol, 2013. 20(11): p. 1364–74.

37. Yang, N., et al., Phosphatidylinositol 4-kinase IIIbeta is required for severe acute respiratory syndrome coronavirus spike-mediated cell entry. J Biol Chem, 2012. 287(11): p. 8457–67.

38. Vella, G., et al., IL-17C contributes to NTHi-induced inflammation and lung damage in experimental COPD and is present in sputum during acute exacerbations. PLoS One, 2021. 16(1): p. e0243484.

39. McGill, J., N. Van Rooijen, and K.L. Legge, IL-15 trans-presentation by pulmonary dendritic cells promotes effector CD8 T cell survival during influenza virus infection. J Exp Med, 2010. 207(3): p. 521–34.

40. Wark, P.A., et al., Diversity in the bronchial epithelial cell response to infection with different rhinovirus strains. Respirology, 2009. 14(2): p. 180–6.

