## Supplementary Material - Results for "PI4KIIIβ inhibition reduces rhinovirus associated cell shedding and ciliary dysfunction"

**Supplementary Material – Figures**

**Supplementary Table 1:** Details of COPD patients and healthy volunteers who participated in the current study

| **Patient ID** | **Age (years)** | **Sex** | **Current Smoker** | **Pack years** | **FEV_1_ actual (L)** | **FEV_1_ %** | **FVC (L)** | **FVC %** | **FEV1/FVC** | **COPD** | **Exacerbations per year** | **Allergies/**  **Hay fever** |
| --- | --- | --- | --- | --- | --- | --- | --- | --- | --- | --- | --- | --- |
| **COPD donors** |  |  |  |  |  |  |  |  |  |  |  |  |
| (1) | 57 | F | No | 39 | 1.25 | 49 | 3.76 | 118 | 0.33 | Yes | >2 | Yes |
| (2) | 71 | F | Yes | 30+ | 1.42 | 66 | 2.13 | 88 | 0.66 | Yes | >2 | Yes |
| (3) | 78 | M | yes | 52 | 1.9 | 22 | 2.6 | 64 | 0.73 | Yes | >2 | No |
| (4) | 76 | F | No | 20 | 1.36 | 40 | 2.6 | 70 | 0.52 | Yes | >2 | No |
| (5) | 77 | F | No | 50 | 0.81 | 45 | 1.76 | 81 | 0.46 | Yes | >2 | No |
| (6) | 74 | M | Yes | 30 | 1.2 | 52 | 2.97 | 97 | 0.4 | Yes | >2 | No |
| (7) | 81 | M | Yes | 75 | 0.41 |  | 1.31 |  | 0.31 | Yes | >2 | No |
| (8) | 75 | M | No | 80 | 1.9 | 75 | 3.3 | 92 | 0.58 | Yes | >2 | No |
| (9) | 61 | M | No | 90 | 1.25 | 40 | 2.54 | 63 | 0.49 | Yes | >2 | No |
| (10) | 75 | F | No | 60 | 0.87 | 46 | 1.48 | 65 | 0.59 | Yes | >2 | No |
| (11) | 71 | M | No | 40 | 0.99 | 37 | 2.77 | 80 | 0.35 | Yes | 1 | Yes |
| **Healthy donors** |  |  |  |  |  |  |  |  |  |  |  |  |
| (12) | 54 | M | No |  |  |  |  |  |  | No |  | Rodents allergy |
| (13) | 59 | M | No |  |  |  |  |  |  | No |  | No |
| (14) | 56 | F | No |  |  |  |  |  |  | No |  | No |
| (15) | 69 | F | No |  |  |  |  |  |  | No |  | No |
| (16) | 60 | M | No |  |  |  |  |  |  | No |  | No |
| (17) | 57 | F | No |  |  |  |  |  |  | No |  | No |
| (18) | 54 | F | No |  |  |  |  |  |  | No |  |  |
| (19) | 54 | F | No |  |  |  |  |  |  | No |  | Yes |
| (20) | 64 | F | No |  |  |  |  |  |  | No |  | No |
| (21) | 62 | F | No |  |  |  |  |  |  | No |  | No |
| (22) | >50 | M | No |  |  |  |  |  |  | No |  | No |
| (23) | 49 | M | No |  |  |  |  |  |  | No |  | No |

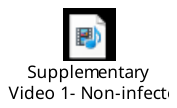

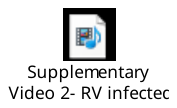

**Supplementary Videos:** High speed videos of non-infected and infected with rhinovirus airway epithelial cultures from a COPD donor at 24hours. Recordings show primary epithelial cells differentiated for 5 weeks at ALI. Supplementary Video 1: Non-infected control represents the airway epithelium covered by ciliated cells beating in a coordinated manner. Supplementary Video 2: Rhinovirus infected airway epithelium shows detached rounded epithelial cells following infection with a few ciliated beating cells remained attached to the epithelial layer.

|  | Healthy | | | COPD | | |
| --- | --- | --- | --- | --- | --- | --- |
| Analyte | **CTRL**  (pg/ml) | **RV**  (pg/ml) | ***P values*** | **CTRL**  (pg/ml) | **RV**  (pg/ml) | ***P values*** |
| IL-1β | 0.79±  0.55 | 7.42±  6.39 | *0.12* | 0.11±  0.01 | 1.24±  0.04 | ***0.016*** |
| IP-10/ CXCL10 | 8.94±  3.19 | 361±  130 | ***9.2e-11*** | 7.27±  1.72 | 555.25±  148.66 | ***4.76e-11*** |
| IL-6 | 359.56±  221.05 | 1193.97±  404.83 | ***0.001*** | 59.22±  18.82 | 300.59±  138.12 | ***0.006*** |
| IL-8/ CXCL8 | 1374.06 ± 217.97 | 2113 ± 105.00 | ***0.016*** | 1330.18 ± 150.11 | 2104.40 ± 108.04 | ***0.045*** |
| RANTES/ CCL2 | 0.88±  0.12 | 7.30±  3.94 | ***3.95e-05*** | 0.96±  0.09 | 5.2±  0.95 | ***0.0001*** |
| TNF-α | 55.10 ± 14.18 | 101.18 ± 22.15 | ***0.003*** | 49.36 ± 6.13 | 97.29 ± 30.18 | *0.052* |
| MCP-1/ CCL2 | 118.11 ± 38.35 | 428.64 ± 125.23 | ***0.00023*** | 136.40 ± 49.69 | 444.14 ± 165.83 | ***0.0030*** |
| ENA-78 | 161.61 ± 67.79 | 390.20 ± 98.94 | *0.14* | 65.73 ± 25.48 | 356.66 ± 190.20 | *0.20* |
| MIP-3α / CCL20 | 102.69 ± 46.20 | 329.15 ± 197.42 | ***0.0018*** | 80.09 ± 21.33 | 190.87 ± 59.27 | ***0.016*** |
| CSF3/ GM-CSF | 23.45±  10.36 | 78.80±  25.56 | ***0.00012*** | 11.92±  0.24 | 21.87±  5.35 | ***0.19*** |
| GRO-α / CXCL1 | 390.15 ± 40.41 | 397.84 ± 22.17 | *0.89* | 401.71 ± 14.08 | 407.07 ± 21.14 | *0.99* |
| IL-15 | 7.85 ± 0.68 | 9.86 ± 0.93 | *0.21* | 7.68 ± 0.55 | 9.30 ± 0.84 | *0.41* |
| IFN-α | 2.56 ± 0.07 | 2.67 ± 0.04 | *0.41* | 2.65 ± 0.06 | 2.80 ± 0.07 | *0.10* |
| IFN-β | 19.15 ± 2.51 | 18.11 ± 3.56 | *0.78* | 16.57 ± 2.14 | 15.70 ± 1.870 | *0.96* |
| IFN-γ | 1.88 ± 0.63 | 1.66 ± 0.38 | *0.99* | 2.05 ± 0.49 | 1.86 ± 0.32 | *0.98* |
| IFN-λ2/3 | 0.46±  0.06 | 208.76±  86.34 | ***4.24e-18*** | 0.46±  0.07 | 283.461±  77.78 | ***4.24e-18*** |
| IL-17c | 65.87±  33.27 | 446.19±  222.00 | ***1.28e-7*** | 28.17±  10.69 | 153.05±  26.76 | ***6.02e-7*** |
| TARC/ CCL17 | 0.22 ± 0.06 | 0.25 ± 0.08 | *0.75* | 0.29 ± 0.06 | 0.36 ± 0.08 | *0.57* |
| CXCL14 | 1175.97 ± 5.52 | 1192.03 ± 8.09 | *0.41* | 1200.04 ± 10.08 | 1207.90 ± 5.83 | *0.78* |
| IL-36g | 11.77 ± 1.00 | 15.27 ± 2.62 | *0.29* | 11.89 ± 0.44 | 13.35 ± 1.16 | *0.71* |

**Supplementary Table 2:** Inflammatory mediators secreted on the basolateral site of airway cultures from healthy and COPD donors at 24hour

Abbreviations: IL-1β = Interleukin-1β; IP-10 = Interferon gamma induced protein 10; IL-6 = Interleukin-6; IL-8 = Interleukin-8; RANTES = regulated on activation, normal T cells expressed and secreted; TNF-α = tumour necrosis factor-α; MCP-1 = monocyte chemoattractant protein-1; ENA-78 = epithelial-derived neutrophil-activating peptide- 78; MIP-3α = macrophage inflammatory protein-3α; GM-CSF = granulocyte colony stimulating factor; GRO-α = growth related oncogene-a; IL-15 = Interleukin- 15; IFN-α = Interferon-a; IFN-β = Interferon-b; IFN-γ = Interferon-g; IFNλ2/3 = Interferon λ 2/3; IL-17c = Interleukin 17c; TARC= TARC - thymus and activation regulated chemokine; CXCL14 – chemokine (C-X-C motif) ligand 14; IL-36g- interleukin 36 gamma

|  | Healthy | | | COPD | | |
| --- | --- | --- | --- | --- | --- | --- |
| Analyte | **CTRL**  (pg/ml) | **RV**  (pg/ml) | ***P values*** | **CTRL**  (pg/ml) | **RV**  (pg/ml) | ***P values*** |
| IL-1β | 0.35±  0.09 | 1.25±  0.55 | *0.17* | 0.32±  0.09 | 2.55±  1.05 | ***0.003*** |
| IP-10/ CXCL10 | 5.94±  1.79 | 177.15±  30.12 | ***2.2E-07*** | 10.55±  3.90 | 246.00±  49.71 | ***2.38E-07*** |
| IL-6 | 110.62±  41.30 | 360.96±  114.66 | *0.57* | 248.54±  134.49 | 872.29±  275.88 | ***0.036*** |
| IL-8/ CXCL8 | 478.78 ± 107.86 | 432.04 ± 86.85 | *0.99* | 376.51 ± 59.73 | 718.25 ± 95.95 | *0.13* |
| RANTES/ CCL2 | 0.43±  0.10 | 6.30±  1.45 | ***0.0008*** | 0.42±  0.09 | 10.62±  4.06 | ***0.0001*** |
| TNF-α | 19.13 ± 4.46 | 25.83 ± 5.22 | *0.79* | 22.62 ± 5.96 | 46.07 ± 11.55 | *0.19* |
| MCP-1/ CCL2 | 42.32 ± 13.94 | 45.57 ± 13.60 | *0.97* | 55.02 ± 21.96 | 145.27 ± 46.40 | *0.29* |
| ENA-78 | 80.64 ± 39.83 | 104.69 ± 30.58 | *0.46* | 188.34 ± 103.67 | 356.20 ± 177.5 | *0.65* |
| MIP-3α / CCL20 | 79.07 ± 50.27 | 46.49 ± 11.49 | *0.98* | 35.08 ± 11.07 | 63.37 ± 19.76 | *0.42* |
| CSF3/ GM-CSF | 7.28±  2.36 | 16.47±  9.35 | *0.44* | 5.70±  1.70 | 15.26±  3.95 | ***0.01*** |
| GRO-α / CXCL1 | 81.51 ± 11.75 | 76.10 ± 11.26 | *0.95* | 63.28 ± 7.82 | 88.03 ± 6.23 | *0.31* |
| IL-15 | 2.60±  0.21 | 2.80±  0.24 | *0.69* | 2.43±  0.16 | 3.57±  0.21 | ***0.005*** |
| IFN-α | 1.06 ± 0.02 | 1.12 ± 0.03 | *0.18* | 1.06 ± 0.01 | 1.27 ± 0.13 | *0.052* |
| IFN-β | 4.37±  0.42 | 6.73±  2.08 | *0.60* | 3.35±  0.37 | 8.13±  1.74 | ***0.01*** |
| IFN-γ | 0.87 ± 0.17 | 0.61 ± 0.09 | *0.75* | 0.82 ± 0.10 | 0.59 ± 0.15 | *0.83* |
| IFN-λ2/3 | 0.19±  0.02 | 196.54±  101.16 | ***1.59E-12*** | 0.44±  0.22 | 300.01±  115.89 | ***9.59E-13*** |
| IL-17c | 1.84±  1.22 | 12.39±  8.45 | *0.09* | 1.31±  0.57 | 13.76±  6.07 | ***0.01*** |
| TARC/ CCL17 | 0.08 ± 0.01 | 0.09 ± 0.02 | *0.98* | 0.07 ± 0.02 | 0.14 ± 0.05 | *0.39* |

**Supplementary Table 3:** Inflammatory mediators secreted in the apical site of airway cultures from healthy and COPD donors at 24hour

Abbreviations: IL-1β = Interleukin-1β; IP-10 = Interferon gamma induced protein 10; IL-6 = Interleukin-6; IL-8 = Interleukin-8; RANTES = regulated on activation, normal T cells expressed and secreted; TNF-α = tumour necrosis factor-α; MCP-1 = monocyte chemoattractant protein-1; ENA-78 = epithelial-derived neutrophil-activating peptide- 78; MIP-3α = macrophage inflammatory protein-3α; GM-CSF = granulocyte colony stimulating factor; GRO-α = growth related oncogene-a; IL-15 = Interleukin- 15; IFN-α = Interferon-a; IFN-β = Interferon-b; IFN-γ = Interferon-g; IFNλ2/3 = Interferon λ 2/3; IL-17c = Interleukin 17c; TARC= TARC - thymus and activation regulated chemokine;

*To note:* CXCL14 and IL-36g were not detected in the apical site of airway cultures.

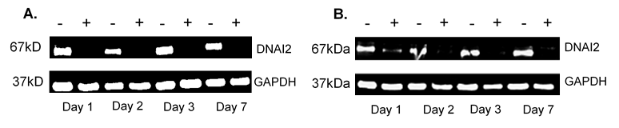

**Supplementary Figure 1:** DNAI2 protein expression during longer term (7 day) rhinovirus infection in airway cultures from healthy and COPD donors

A) Representative images of DNAI2 expression in cultures from healthy individuals;

B) Representative images of DNAI2 expression in cultures from COPD individuals; ‘-‘ depicts mock-infected and ‘+’ depicts rhinovirus infected

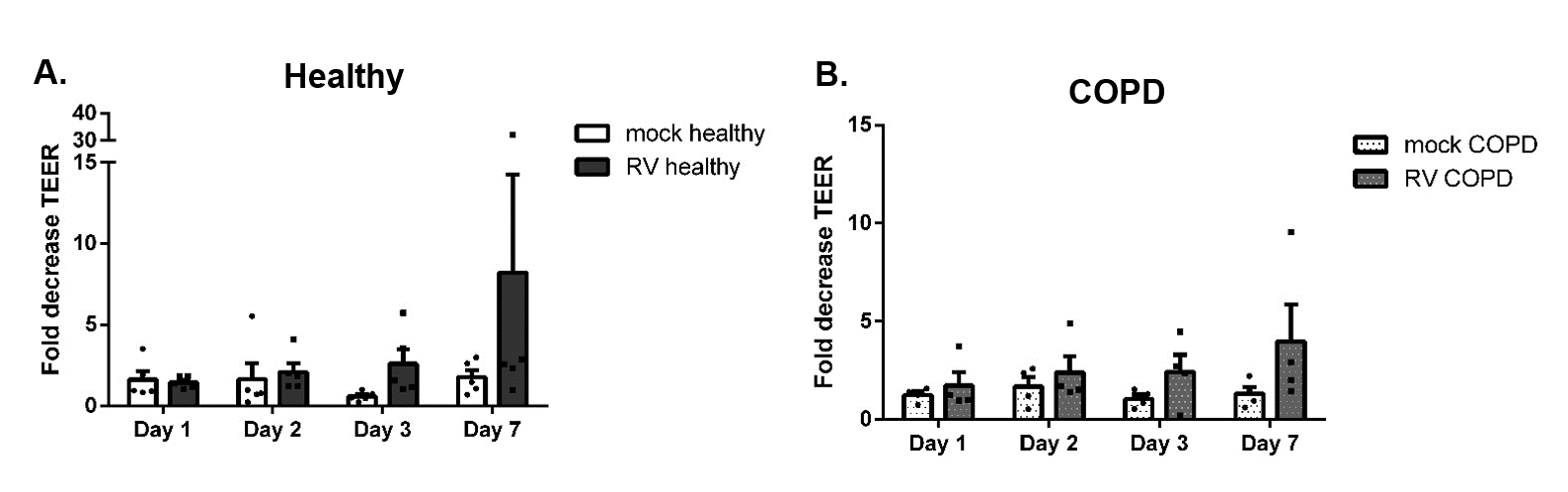

**Supplementary Figure 2:** Permeability and integrity of airway ciliated cultures during longer term (7 day) rhinovirus infection

A) Transepithelial electrical resistance (TEER) of mock and rhinovirus infected airway cultures from A) healthy individuals and B) COPD at days 1, 2, 3 and 7 expressed as fold change relative to day 0 measurements; n=2 in 4 donors/ group.

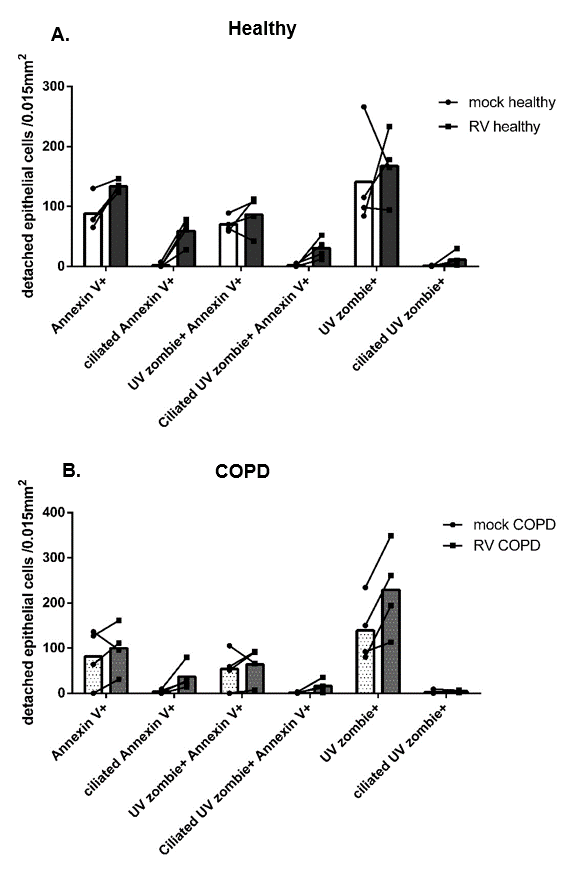

Supplementary Figure 3: Epithelial cells detached from healthy and COPD epithelium infected with rhinovirus for 24hour.

A) Number of detached epithelial cells in healthy apical secretions positive for Annexin V, UV zombie dye or Annexin V and UV zombie dye post rhinovirus infection counted per 0.015mm^2^ field from 25 random images from an apical culture washing; B) Number of detached epithelial cells in COPD apical secretions positive for Annexin V, UV zombie dye or Annexin V and UV zombie dye post rhinovirus infection counted per 0.015mm^2^ field from 25 random images from an apical culture washing. Apoptotic cells are positive for Annexin V marker, late apoptotic cells are positive for Annexin V and UV zombie markers, necrotic/ dead cells are positive for UV zombie dye; n=4 donors/ group
