## Supplementary Material - Methods for "PI4KIIIβ inhibition reduces rhinovirus associated cell shedding and ciliary dysfunction"

**Collection of primary human respiratory epithelial cells**

Primary airway epithelial cells were collected by brushing the interior part of the nasal turbinate of healthy or COPD donors as previously described [1]. Cells were resuspended in 20mM HEPES buffered medium199 (Invitrogen) with 100IU/ml penicillin, 100 μg/ml streptomycin (Invitrogen), 50 μg/ml gentamicin and 2.5 μg/ml fungizone (Invitrogen) [2]. Ethical approval for sample collection was obtained through the UCL Living Airway Biobank (Ref: 14/NW/0128).

**Culture of primary human respiratory epithelial cells**

Co-culture of human airway cells with mitotically inactivated feeder layers was carried out as previously described [3]. Human primary cells were fed every other day with 3:1 DMEM with 10% foetal bovine serum (Invitrogen), 1% penicillin/streptomycin and F-12 supplemented nutrient mix (Invitrogen) and 5μM Y-27632 (Cambridge Bioscience, UK) [2]. Human primary cells were cultured at 4x10^5^cells/0.33cm^2^ transwell insert (Corning, USA) in submerged conditions for 2 days at 37^o^C and 5%CO_2_. Thereafter, primary cells were grown at air-liquid interface (ALI) in 1:1 DMEM:airway epithelial cell growth medium containing the supplement kit provided by the manufacturer (AECGM, Promocell, Germany), then supplemented with 100nM retinoic acid for 28 days [1, 2].

**TEER measurement**

TEER was determined using an EVOM2 voltmeter and a pair of STX2 chopstick electrodes (World Precision Instruments, USA). The meter reading of an empty insert was used to set a baseline value. All readings were in a digital format which provided a range of 1-9,999 Ohms (Ω). To measure TEER of ALI cultures, 0.2ml of BEBM medium (Lonza, UK) was added to the apical surface where the electrodes were placed for 2 seconds. After the measurement was taken the BEBM medium was removed from the apical surface.

**Virus culture**

Rhinovirus 16 was kindly provided by Prof Gary McLean (Cellular and Molecular Immunology Research Centre, London, UK) was propagated in Hela H1 cells also provided by Prof Gary McLean. Briefly, 5ml of rhinovirus stock made up in 2% Foetal Calf Serum, 1% Penicillin/Streptomycin, 15mM HEPES (Sigma-Aldrich) and 15mM Sodium bicarbonate (Sigma-Aldrich) in DMEM media was expanded to 25ml through passaging in cell culture for 24h until >80% cytopathic effects were observed [4]. Infected cell lysates were clarified through centrifugation 4000 x *g* for 15min and filtered through 0.2μm vacuum filters, followed by centrifugation in 100 kDa NMCO Amicon ultra devices (AmiconUltra, Merck, U centrifugal filtration) at 3000 x *g* at 4^o^C. Virus was precipitated using 7% polyethylene glycol 6000 (Sigma-Aldrich) and 0.5M sodium chloride (Sigma-Aldrich). Titre of the virus was determined by titration assays in Hela H1 cells and TCID_50_ was calculated using the Spearman-Karber equation [5].

**Rhinovirus *in vitro* infection of human airway ciliated cultures**

An aliquot of 1.5x10^6^ TCID_50_/ml from the viral stock was added to the apical site of epithelial cultures in 100μl BEBM media (Lonza, UK) .Virus was allowed to attach for 1hour at 37^o^C with 5%CO_2_ and was then removed to allow infection progression for further 24hour or 7 days at air-fluid interface. Mock infected cultures received 100μl of BEBM (Lonza, UK) media and were processed as infected wells.

**PI4KIIIβ** **inhibition of rhinovirus replication in airway epithelial cells**

Epithelial cultures were pre-treated with a PI4KIIIβ inhibitor (GSK’533) [6] supplied by the Refractory, Respiratory and Inflammation Unit at GSK, Stevenage at a final concentration of 100nM added to the basolateral medium for 1h at 37^o^C and 5% CO2. Vehicle control received DMSO to the basolateral medium, and was prepared in the same manner as the inhibitor compound where DMSO final concentration was lower than 0.5% and further diluted in assay medium. Cultures were then infected with 1.5x10^6^ TCID50/ml apically in 100μl of BEBM media for 1h at 37^o^C with 5% CO_2_. Mock infected controls received 100μl BEBM media only and were treated in the same manner as infected wells. Infection was allowed to progress for further 24hours at 37^o^C with 5% CO_2_ at air-fluid interface.

**Ciliary function analyses**

Recordings of cilia beating was carried out by a Hamamatsu ORCA digital camera C11440 (Hamamatsu, Japan) attached to an inverted phase-contrast Nikon Eclipse Ti-E (Nikon instruments, UK) microscope and a 20x long distance objective at 394 frames per second (fps) at 512 x 512 pixel resolution with 2ms exposure time. Ciliary beat frequency (CBF) was determined as described previously [7]. Ciliary beat activity was calculated by CiliaFa software [8].

**Immunofluorescence microscopy**

Epithelial cells were fixed with 4% PFA for 20min at room temperature. Staining procedure in transwell inserts was carried as previously described [2, 7]. Primary antibodies anti-VP2 (1:200, QED biosciences, USA), anti-β tubulin (Abcam, UK), anti-PI4P (1:500, Echelon Biosciences, UK) were detected by anti-rabbit A488 (1:1000, Abcam, UK), anti-mouse A647 (1:1000, Biorad, UK) and anti-mouse A488 (Abcam, UK). Anti-VP2 antibody was also conjugated to A647 fluorophore (1:200, Abcam, UK) to avoid host species cross-reactivity. Hoechst staining solution 33258 (Sigma-Aldrich, UK) was applied apically for 30min to counterstain cell nuclei. Membranes with epithelial cells were mounted onto glass slides with n-propylgallate and imaged by Zeiss LSM 710 confocal microscope (Carl Zeiss ltd, Germany).

**Western blot analysis**

Cell lysates were prepared in RIPA buffer (Sigma-Aldrich) containing 1% protease inhibitors cocktail (Complete, Roche). Protein concentration was determined by BCA assay (Pierce, Thermo-fisher). Samples were separated by Sodium Dodecyl Sulphate-Polyacrylamide Gel Electrophoresis (SDS-PAGE) using 12% acrylamind gels and electro-transferred onto Polyvinylidene fluoride or polyvinylidene difluoride (PVDF) membranes. Membranes were probed for human anti-DNAI2 (1:500, mouse, Bio-Techne) and anti-GAPDH (1:2000, rabbit, Abcam) at 4^o^C overnight, followed by incubation with anti-mouse (1:1000, Biо-techne) and anti-rabbit (1:1000, Cell signalling technologies) horseradish peroxidase linked antibodies for 1h. Blots were finally washed 4 times in PBST (0.05% Tween-20) and stained with enhanced chemiluminescence (ECL) substrate (BioRad, UK). Densitometry analysis was performed using ImageJ software 1.51j8 (National Institutes of Health, USA). To determine the abundance of the target proteins, percent values were normalised to relative GAPDH.

**Apoptosis analysis**

Epithelial cells detached from the epithelial layer post infection were stained with UV zombie dye (1:500, BioLegend) for 15min in Annexin V Binding Buffer (0.1M HEPES, 1.4M NaCl and 25mM CaCl_2_) (BD Pharmingen). Cells were washed in Annexin V Binding Buffer, centrifuged at 3700 x *g* for 5min and stained with 5μl/sample Annexin V dye (BD Pharmingen) for 15min, followed by another wash. Cell pellets were fixed in 1% PFA for 20min and then cytospun at 120 x *g* for 5min onto glass slides and mounted with N- propylgallate (Sigma-Aldrich). Images were taken on a Nikon TiE-Eclipse 100 microscope.

**Cytokine and chemokine measurement**

Cytokines and chemokines were measured using 96-well MSD plates (Meso Scale, UK) and Luminex plex plates (Invitrogen, UK) according to manufacturer’s instructions. The pro-inflammatory mediators IL-1β, IP-10/ CXCL10, IL-6, IL-8/ CXCL8, TNF-α, RANTES/ CCL5, MCP-1/ CCL2, ENA-78/ CXCL5, MIP-3α/ CCL20, GRO-α/ CXCL1, CSF-3/ GM-CSF, IL-15 were quantified using a custom 10 spot Luminex plex. Plates were read by a Bio-Plex 3D Suspension Array instrument (Bio-Rad, UK). The concentration of the samples was calculated by plotting the expected standards concentration against their mean fluorescence intensity. The interferons IFN-α, IFN-β, IFN-λ, IFN-γ were quantified by U-plex plates and the pro-inflammatory cytokines IL-17c and TARC/ CCL17 were quantified by V-plex plates using a MSD plate reader Sector Imager 3000 (MSD, UK). Data were calculated by the Discovery Workbench software version 3.0 (MSD) by plotting the standard curves of calibrators and samples absorbance readings against the relevant standard curves.

**Statistical analyses**

Statistical analyses were performed using the statistical programming language R (R-core team 2018), Version 3.5.1. For experiments involving different treatments or time points, data was pooled for all experimental conditions and relevant disease groups were analysed jointly using linear modelling techniques. Technical replicates were averaged prior to modelling. All models included separate means (*e*.*g.* fixed effects) per experimental condition and disease group, with included random effect for each donor to acknowledge donor dependence. For studies where only one observation per donor was collected, standard linear regression models were fitted. Multiple comparison correction using the Benjamini-Hochberg (BH) method was applied to all p-values. Paired t-test were applied for experiments with only two experimental conditions where differences for each disease group were of interest. Graphs were plotted using Prism v6.1 (GraphPad Software, USA).

For multiple cytokines and chemokines analysis, all values were normalised by log transformation. Any missing values were imputed with a lower detection limit (LLOD) calculated from MSD Workbench or the lowest detected value. Principal component analysis (PCA) was conducted in R studio (v1.4).

1.	Hirst, R.A., et al., *Ciliated air-liquid cultures as an aid to diagnostic testing of primary ciliary dyskinesia.* Chest, 2010. **138**(6): p. 1441-7.

2.	Lee, D.D.H., et al., *Ciliated Epithelial Cell Differentiation at Air-Liquid Interface Using Commercially Available Culture Media.* Methods Mol Biol, 2020. **2109**: p. 275-291.

3.	Butler, C.R., et al., *Rapid Expansion of Human Epithelial Stem Cells Suitable for Airway Tissue Engineering.* Am J Respir Crit Care Med, 2016. **194**(2): p. 156-68.

4.	Walker, E.J., et al., *Rhinovirus 3C protease facilitates specific nucleoporin cleavage and mislocalisation of nuclear proteins in infected host cells.* PLoS One, 2013. **8**(8): p. e71316.

5.	Papi, A. and S.L. Johnston, *Rhinovirus infection induces expression of its own receptor intercellular adhesion molecule 1 (ICAM-1) via increased NF-kappaB-mediated transcription.* J Biol Chem, 1999. **274**(14): p. 9707-20.

6.	Roulin, P.S., et al., *Rhinovirus uses a phosphatidylinositol 4-phosphate/cholesterol counter-current for the formation of replication compartments at the ER-Golgi interface.* Cell Host Microbe, 2014. **16**(5): p. 677-90.

7.	Smith, C.M., et al., *Ciliary dyskinesia is an early feature of respiratory syncytial virus infection.* Eur Respir J, 2014. **43**(2): p. 485-96.

8.	Smith, C.M., et al., *ciliaFA: a research tool for automated, high-throughput measurement of ciliary beat frequency using freely available software.* Cilia, 2012. **1**: p. 14.
